# SSNA1 mechanically reinforces the damaged microtubule lattice

**DOI:** 10.64898/2026.01.16.699973

**Authors:** Laura B. Richardson, Elizabeth J. Lawrence, Abinaya Pinjakan, Marija Zanic

**Author notes:** Correspondence to: Marija Zanic. **Author Contributions** LBR, EJL, and MZ designed research; EJL and LBR generated reagents; LBR performed experiments; LBR and AP analyzed data; LBR and MZ wrote the paper.

## Abstract

SSNA1 (Sjögren’s Syndrome Nuclear Autoantigen 1) is a microtubule-associated protein involved in key cellular processes, including cell division, intraflagellar transport, and axonal branching. SSNA1 specifically localizes to sites of damage along the microtubule lattice, thus acting as a microtubule damage sensor. However, the effects of SSNA1 on microtubule mechanics or on the process of microtubule self-repair, which involves the incorporation of soluble tubulin dimers into lattice damage sites, are not known. Here, we use *in vitro* reconstitution with purified proteins and total internal reflection fluorescence (TIRF) microscopy to probe SSNA1’s effects on microtubule mechanics and self-repair. We apply two distinct sources of force to investigate microtubule mechanics: kinesin-driven gliding assays and microfluidic flow. We find that SSNA1 binding increases microtubule rigidity and resistance to breakage under the physiological and controlled forces in our assays. Interestingly, SSNA1’s localization to microtubule damage sites prevents the incorporation of new tubulin dimers and thus inhibits lattice self-repair. Conversely, we find that SSNA1 does not recognize damage sites that have been repaired by tubulin incorporation. Together, our findings demonstrate that SSNA1 reinforces the mechanical strength of microtubules without promoting self-repair, suggesting an alternative mechanism for restoring microtubule integrity in the absence of tubulin-mediated repair and providing new insights into SSNA1’s mechanism of microtubule stabilization.

**Significance Statement:** Microtubules are cytoskeletal polymers that experience mechanical stress during essential cellular processes such as cargo transport, cell division, and ciliary beating. To maintain their integrity, microtubules rely on both stabilizing proteins and repair mechanisms. Here, we show that microtubule-associated protein SSNA1 strengthens microtubules by increasing their rigidity and resistance to force-induced breakage, while simultaneously blocking tubulin-mediated lattice repair at sites of damage. By distinguishing between damaged and repaired microtubule lattices, SSNA1 enforces a stabilization strategy that favors mechanical reinforcement over self-repair. These findings reveal a new mode of microtubule regulation that decouples mechanical stability from lattice repair and provide insight into how cells preserve cytoskeletal integrity under force.

## Introduction

Microtubules are dynamic cytoskeletal polymers that play crucial mechanical roles within the cell (Alberts 2022). As the most rigid component of the cytoskeleton, microtubules provide structural backbones for cellular morphologies ranging from motile cilia and flagella to elongated axons (Baas et al. 1988; Gittes et al. 1993; Mickey and Howard 1995; Satir and Christensen 2007; Howard 2001). They withstand compressive and tensile forces in cells like cardiomyocytes and modulate cardiac mechanics (Caporizzo and Prosser 2022; Nishimura et al. 2006). Furthermore, through their controlled polymerization and depolymerization, microtubules themselves generate pushing and pulling forces essential for cellular processes such as mitotic spindle assembly, spindle positioning, and proper chromosome segregation (Dogterom and Yurke 1997; Koshland et al. 1988; McIntosh et al. 2008; Pavin and Tolić 2021; Inoue’t and Salmont, n.d.; Grishchuk et al. 2005). All these cellular functions are supported by the mechanical properties of individual microtubules, which are a direct consequence of their tubular multi-protofilament structure. While cytoskeletal filaments in general exhibit material rigidity comparable to those of hard plastics, the typical 13-protofilament, cylindrical microtubule lattice has a significantly higher bending stiffness when compared to other cytoskeletal polymers due to its larger cross-sectional area (Brangwynne et al. 2006; Gittes et al. 1993; Mickey and Howard 1995; Venier et al. 1994; Hawkins et al. 2010; Howard 2001).

The integrity of the microtubule lattice is closely intertwined with its mechanics. Despite the strong longitudinal and lateral bonds between tubulin dimers that comprise the microtubule polymer, the polymer lattice can incur loss of tubulin dimers through various cellular processes (Dye et al. 1992; Schaedel et al. 2015; Verhey and Ohi 2023; Théry and Blanchoin 2021). In addition to specialized severing enzymes, which use the energy from ATP hydrolysis to extract tubulin dimers out of the lattice (Vemu et al. 2018; Roll-Mecak and McNally 2010), kinesin motor proteins have also been recently reported to induce damage as they translocate along the microtubule (Andreu-Carbó et al. 2022; Budaitis et al. 2022; Triclin et al. 2021). Exertion of successive mechanical force to create damage sites at which tubulin subunits are expelled from the lattice has been shown to soften the microtubule polymer (Memet et al. 2018; Schaedel et al. 2015). Importantly, such lattice damage sites can be ‘repaired’ through reincorporation of new tubulin dimers from solution into the polymer lattice (Gazzola et al. 2022; Schaedel et al. 2015). And while microtubule polymer damage can weaken microtubule mechanical strength and lead to the premature release of motor proteins (Liang et al. 2016), the process of microtubule self-repair results in overall increased polymer stability and longevity. Specifically, microtubule self-repair generates new regions of the microtubule lattice that can withstand depolymerization, promote transitions from microtubule shrinkage back to growth, and even protect the lattice against further damage (Aumeier et al. 2016; de Forges et al. 2016; Rai et al. 2021; Schaedel et al. 2019; Schaer et al. 2023; Triclin et al. 2021; Vemu et al. 2018). Furthermore, recent studies have implicated several microtubule-associated proteins (MAPs) in the process of microtubule repair (de Forges et al. 2016; Aher et al. 2020; Duan et al. 2023; Van Den Berg et al. 2023; Biswas et al. 2024).

SSNA1 (Sjögren’s Syndrome Nuclear Autoantigen 1) is a MAP recently shown to act as a sensor of microtubule lattice damage *in vitro* (Lawrence et al. 2021). SSNA1 is a small, ∼14 kDa protein that self-assembles into fibrils (Basnet et al. 2018; Ramos-Morales et al. 1998; Rodríguez-Rodríguez et al. 2011). It localizes to basal bodies of cilia and flagella, centrosomes in dividing cells, and axonal branch points in neurons, and plays important roles in ciliogenesis, cell division and neuronal development (Basnet et al. 2018; Errico 2004; Goyal et al. 2014; Pfannenschmid et al. 2003; Ramos-Morales et al. 1998; Lai et al. 2011). In vitro, SSNA1 stabilizes dynamic microtubule ends and detects microtubule lattice damage (Lawrence et al. 2021). Although SSNA1 localizes to sites of increased microtubule mechanical strain in cells and detects microtubule lattice damage in vitro, the effects of SSNA1 on microtubule polymer mechanics and the process of microtubule lattice repair are not known. Here, we show that SSNA1 mechanically reinforces damaged microtubule lattice but does not promote tubulin-mediated microtubule repair. Rather, SSNA1’s localization limits tubulin incorporation into the sites of lattice damage. Furthermore, SSNA1 does not recognize damage sites that have been repaired by tubulin. Our findings suggest an alternative mechanism for restoring microtubule integrity in the absence of tubulin-mediated repair. Taken together, these results provide new insights into the processes of microtubule lattice damage sensing and repair, and their effects on overall microtubule polymer mechanics.

## Results

### SSNA1 reduces microtubule curvature and protects against microtubule rupture under physiological force

To probe the direct effects of SSNA1 on microtubule mechanics without the confounding variables present in the cell, we employed a reductionist *in vitro* reconstitution approach with purified protein components. To this end, we performed microtubule gliding assays in which purified kinesin motors were bound to coverslip surfaces, and fluorescently labeled Taxol-stabilized microtubules were propelled by the motors and visualized by total internal reflection fluorescence (TIRF) microscopy (Fig. 1A) (see Methods). First, microtubules were introduced into the flow chamber and allowed to bind to kinesin-coated surface in the absence of ATP. Next, the flow chamber was incubated with 500 nM SSNA1 for 20 minutes, resulting in binding of SSNA1 to the microtubule lattice. Excess SSNA1 was washed out of solution, and more microtubules were added into the flow chamber. Because of SSNA1’s low turnover rate on microtubules (Lawrence et al. 2021), this procedure resulted in two populations of microtubules: one coated and one uncoated with SSNA1. After the addition of ATP, microtubules were propelled across the coverslip surface by kinesin motors and both the microtubule and SSNA1 signal were imaged over time (Fig. 1B). This assay allowed us to assess how SSNA1 affects microtubule rigidity under conditions of physiological levels of force through analyzing the extent of microtubule curvature generated by kinesin-driven gliding.

**Figure 1.**
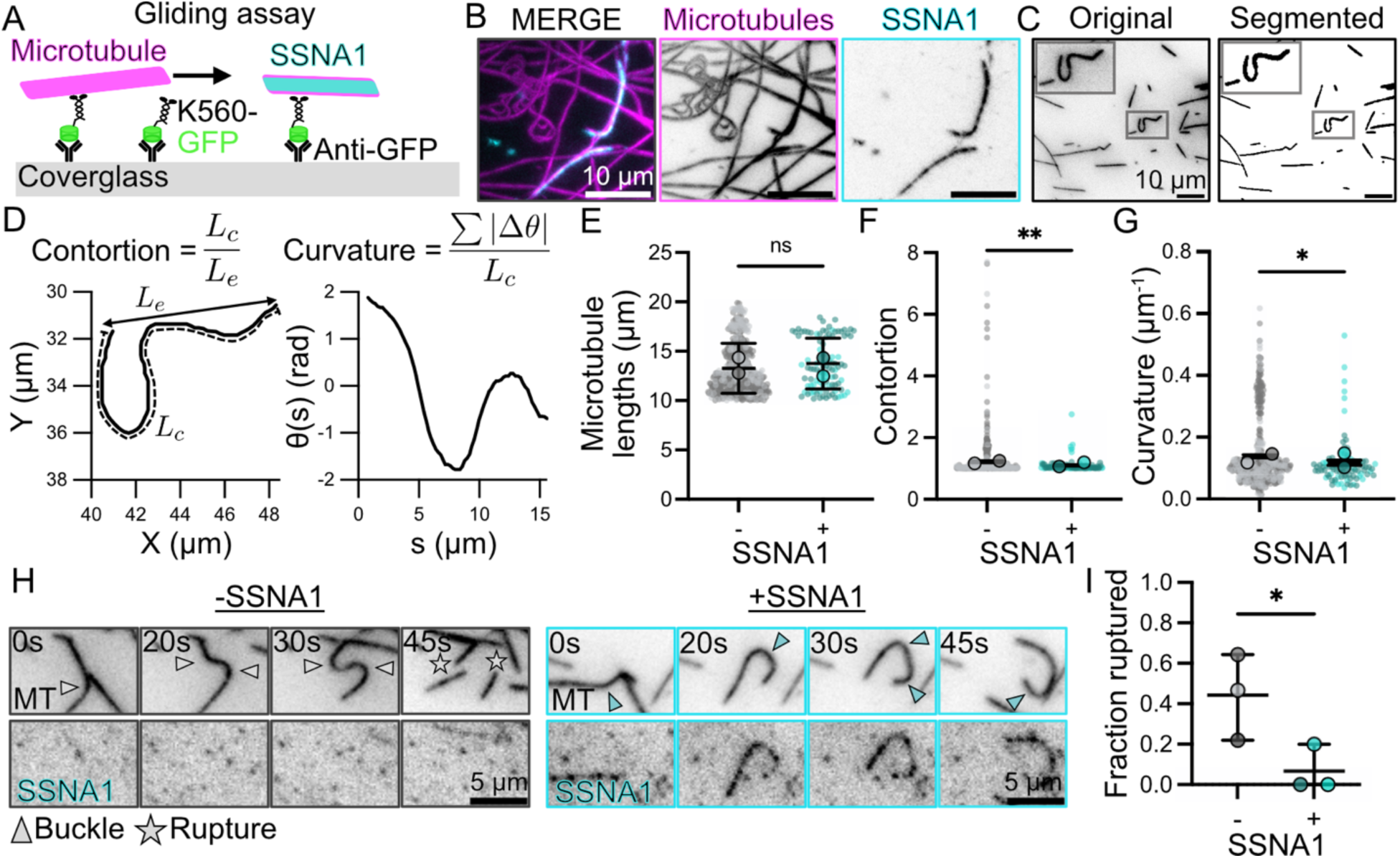
SSNA1 increases microtubule rigidity and mechanical integrity under physiological force. (**A)** Experimental setup schematic. (**B)** Max-intensity projection over 5 minutes of microtubule gliding assay with a mixed population of SSNA1-coated and control microtubules. (**C)** Representative images of Ilastik image segmentation. (**D)** LEFT Representative plot of microtubule coordinates from image segmentation of the microtubule shown in zoom-in window in (C). Contour length is measured along the length of the microtubule and end-to-end distance is defined as the shortest distance between endpoints. RIGHT Representative plot of the change in angle of the example microtubule across its arc length. **(E)** Length-matched microtubule populations used for analysis of curvature. Individual points represent single microtubules. Different shades represent separate experimental days. Lines with error bars show mean ± standard deviation. p = 0.1 (Welch’s t-test). **(F)** Contortion of control and SSNA1-coated microtubules. p = 0.002 (Kolmogorov-Smirnov test). Lines with error bars show mean ± standard error of the mean. (**G)** Curvature of control and SSNA1-coated microtubules. Lines with error bars show mean ± standard error of the mean. p = 0.03 (Welch’s t-test). (E-G): n = 482 microtubules (CTRL), 93 microtubules (+SSNA1); N = 2 experimental days. (**H)** Representative images of GMPCPP-stabilized control and SSNA1-coated microtubules buckling and breaking under the force of kinesin propulsion over time. Arrows: buckling points, stars: microtubule rupture events. (**I)** Fraction of buckled control and SSNA1-coated microtubules that rupture. Shading represents different experimental days. Lines and error bars show mean ± standard deviation. P = 0.04 (paired t-test). n = 38 buckled microtubules (CTRL), 13 buckled microtubules (+SSNA1); N = 3 experimental days.

To measure the individual polymer curvature, microtubule images were segmented using Ilastik machine learning software (Fig. 1C) and sorted into control (-SSNA1) and +SSNA1 categories based on their SSNA1 intensity (see Methods). The cartesian coordinates of segmented and skeletonized microtubules were used to calculate the extension of the polymer based on the ratio of microtubule contour length to end-to-end distance, as well as the average local curvature of the polymer (Fig. 1D, see Methods). To compare polymers of consistent lengths, we only considered microtubules between 10 and 20 microns and confirmed that the length distributions of the two populations were not statistically significantly different (Fig. 1E). Our curvature analysis revealed that the uncoated control microtubules were significantly more contorted than those coated with SSNA1 (1.22 ± 0.03 mean ± SEM, n = 482 control microtubules vs 1.10 ± 0.02, n = 93 for SSNA1-coated microtubules) (Fig. 1B, 1F). Furthermore, control microtubules displayed a significantly higher mean local curvature than those coated with SSNA1 (0.138 ± 0.005 µm^-1^ mean ± SEM, n = 482 control microtubules vs 0.117 ± 0.008 µm^-1^, n = 93 for SSNA1-coated microtubules) (Fig 1G). The lower curvature of SSNA1-coated microtubules suggested that SSNA1 binding increases the bending rigidity of the polymer.

In our gliding assays, we observed many microtubules that buckled under the propulsive force of the surface-attached kinesins. We wondered how binding of SSNA1 may alter the propensity of microtubules to rupture due to buckling. Reported incidence of microtubule rupture for Taxol-stabilized microtubules in gliding assays is exceedingly low (Tsitkov et al. 2022), a finding that was consistent in our experiments. However, when instead of Taxol-stabilized, we introduced GMPCPP-stabilized microtubules in our gliding assay, we observed a measurable frequency of microtubule rupture upon buckling. We therefore specifically focused on the subset of microtubules that exhibited buckling behavior and determined the fraction of buckled microtubules that ruptured in both control and SSNA1-coated populations. We found that SSNA1-coated microtubules had a significantly reduced fraction of ruptured microtubules (7.7%, 1 ruptured out of 13 buckling events) when compared to the control population (47%, 18 ruptured out of 38 buckled microtubules) (Fig. 1H, 1I). Taken together, the results from our gliding assays demonstrate that SSNA1 increases microtubule rigidity and protects against microtubule rupture under physiological force.

### SSNA1 stiffens the microtubule lattice through localization to sites of mechanical damage

The finding that SSNA1 significantly reduced microtubule curvature in gliding assays prompted us to further investigate SSNA1’s effects on microtubule mechanics under more controlled conditions. To that end, we employed a microfluidics-based approach (Rogers et al. 2025) to apply a controlled, uniform force to all microtubules within a field of view in the presence and absence of SSNA1. We attached GMPCPP-stabilized microtubule “seeds” to the surface of our PDMS-fabricated microfluidics device, from which we polymerized microtubule extensions with soluble tubulin and GTP (see Methods). After extensions reached the desired length, we introduced 2 µM tubulin and GMPCPP to “cap” the microtubule extensions and prevent depolymerization (Fig 2A). Since microtubule seeds were stably attached to the surface of the coverslip, this setup allowed us to bend microtubule extensions under controlled hydrodynamic flow without microtubules floating away in solution.

**Figure 2.**
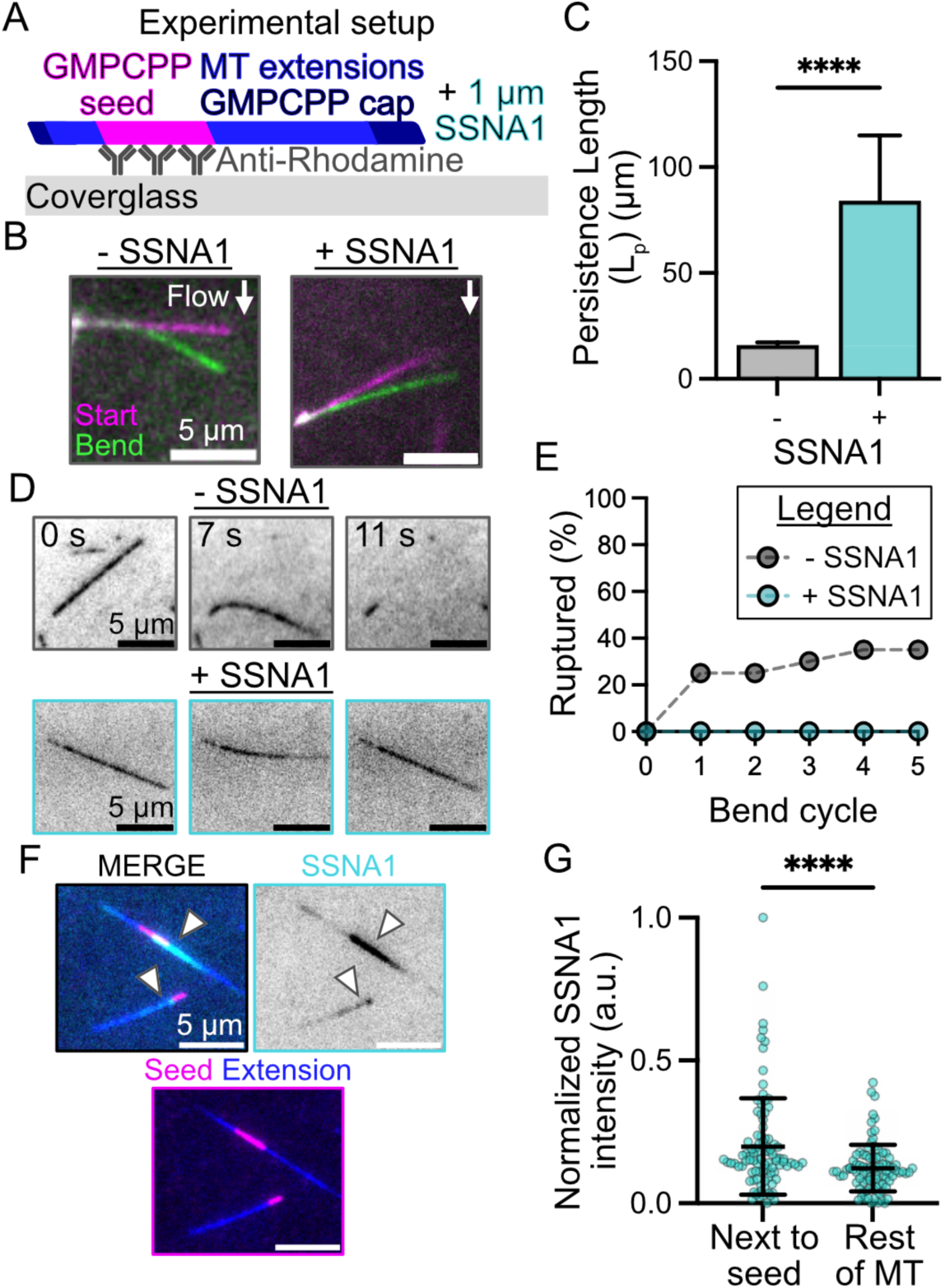
SSNA1 stiffens the microtubule lattice through localization to sites of mechanical damage. **A** Experimental setup schematic. GMPCPP seeds are anchored to the coverslip with capped extensions free to bend. **B** Representative overlay of microtubules bending under one flow pulse. **C** Persistence lengths of microtubules ± SSNA1. Lines with error bars represent mean ± standard deviation. p = <0.0001 (Welch’s t-test) n = 62 microtubules (CTRL), n = 59 microtubules (+SSNA1); N = 2 experimental days. **D** Representative images of microtubule extensions ± SSNA1 buckling and breaking under hydrodynamic force. **E** Fraction of buckled microtubules ± SSNA1 that rupture. n = 20 microtubules (CTRL), 20 microtubules (+SSNA1); N = 2 experimental days. **F** SSNA1 localization on microtubules subjected to pulsatile microfluidic flow. **G** SSNA1 intensity at regions within 2 µm of the adhered microtubule seed vs the rest of the extension. p= <0.0001 (paired t-test). n = 94 microtubules.

We cyclically bent microtubules that had either been pre-incubated with 1 µM SSNA1 or untreated (control microtubules) using cycles of pulsatile flow in our microfluidic device. We observed that the relative displacement of microtubule extensions under flow was larger for control microtubules than those coated with SSNA1, indicating that SSNA1 localization increased the resistance to bending (Fig. 2B). Indeed, our calculations showed a significant increase in microtubule persistence length in the SSNA1-coated microtubule population when compared to the untreated control (Lp = 84.1 ± 30.8 µm, n = 59 SSNA1-coated microtubules, vs. Lp = 16.0 ± 1.3 µm, n = 62 control microtubules) (Fig. 2C). Furthermore, consistent with our gliding assay results, we observed a stark increase in the number of control microtubules that ruptured in the microfluidics assay, with 40% of control microtubules rupturing (8 ruptured out of 20 bent), compared to no rupture events for microtubules coated with SSNA1 (0 ruptured out of 20 bent), even after five cycles of applied hydrodynamic force (Fig. 2D-E). These results demonstrated that the presence of SSNA1 increases microtubule bending stiffness and mechanically reinforces microtubules subjected to controlled flow forces.

To determine whether SSNA1’s effects on microtubule mechanics are a consequence of uniform localization along the microtubule lattice or recognition of specific microtubule features, we imaged fluorescent SSNA1 localization in the microfluidics assay. Our analysis showed that SSNA1 preferentially bound to microtubule extension areas adjacent to the surface-attached microtubule seeds, with significantly higher average SSNA1 signal within 2 μm regions on either side of the microtubule seed as compared to the rest of the extension (Fig. 2F-G). This result is consistent with our previous findings that SSNA1 specifically recognizes sites of microtubule damage (Lawrence et al. 2021), as we expect that flow-induced bending of microtubules preferentially damages microtubule lattice at points of high mechanical strain at the interface of the attached seed and the free microtubule extension. Taken together, our results suggest that SSNA1 mechanically reinforces the microtubule lattice through its specific binding to microtubule damage sites.

### SSNA1’s recognition of microtubule damage limits tubulin-mediated repair

SSNA1’s preferential binding to microtubule damage sites led us to question if SSNA1 may be promoting microtubule self-repair via tubulin incorporation, as has been reported for several other microtubule-associated proteins (de Forges et al. 2016; Aher et al. 2020; Duan et al. 2023; Van Den Berg et al. 2023). To study microtubule self-repair, we first induced damage to Taxol-stabilized microtubules by transiently removing Taxol from solution (Figure 3A, see Methods). This treatment resulted in damaged regions with a broad range of damage sizes along the microtubule lattice, as quantified using a custom-developed damage-detection MATLAB code (see Methods). We classified the damage sites into three size categories (Fig. 3C; <0.5 µm, 0.5 – 1 µm, 1 – 3 µm), which allowed us to investigate potential trends across different damage sizes. We then compared tubulin localization to the damaged microtubule lattices in the absence of SSNA1 to those pre-incubated with SSNA1 (Fig. 3B), ensuring that the sizes of the investigated damaged sites were not significantly different between the two conditions (Fig. 3C).

**Figure 3.**
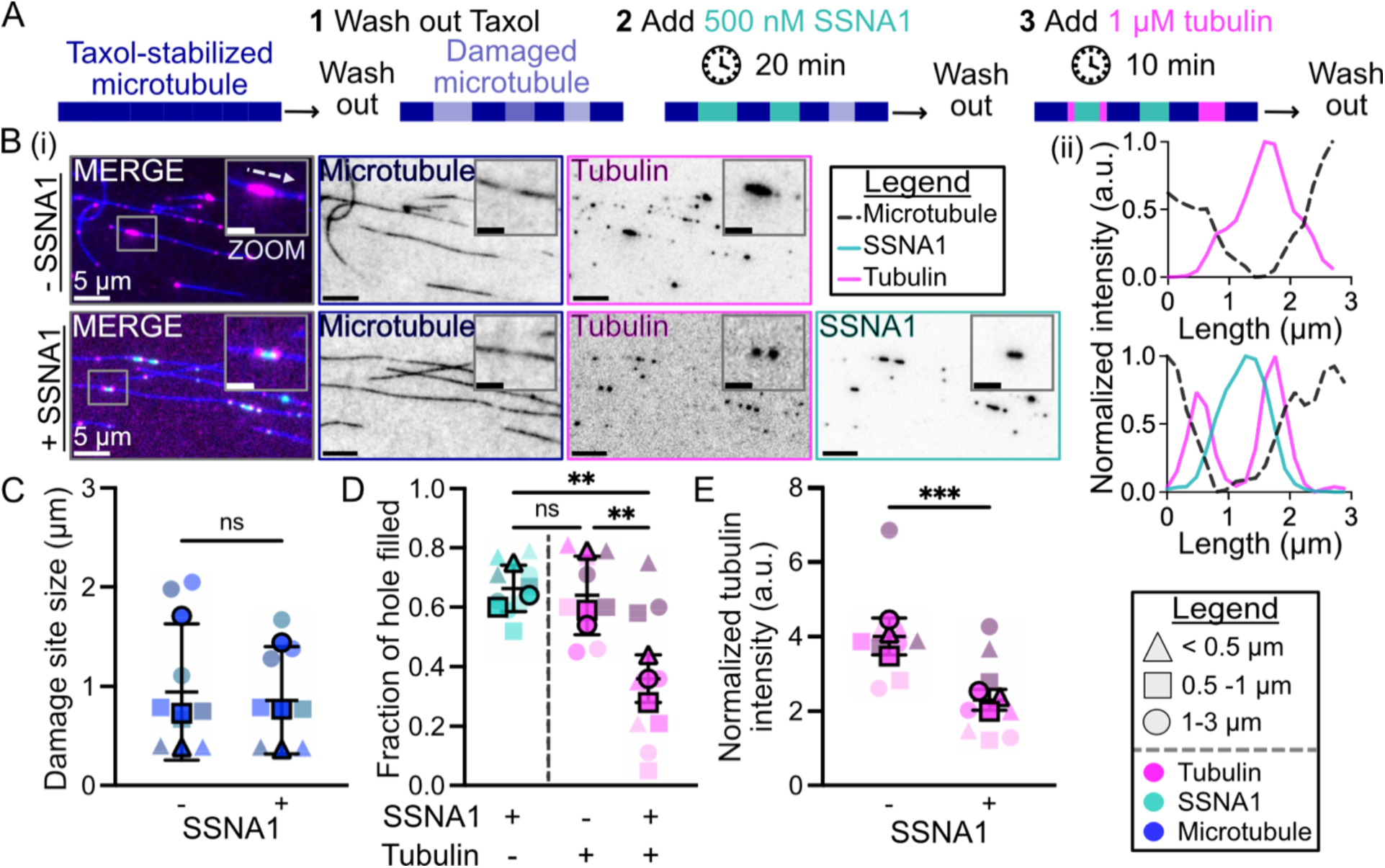
SSNA1 limits tubulin incorporation into damage sites. **(A)** Experimental setup schematic. (**B)** (i) Representative images of microtubule repair assay on damaged microtubules in the absence and presence of SSNA1. Zoom scale bars represent 2 µm. (ii) Representative linescan of tubulin and SSNA1 intensities across a microtubule damage site. **(C)** Distribution of damage site sizes detected during analysis. p = 0.54 (paired t-test). (**D)** Comparison of SSNA1 and tubulin incorporation at damage sites. SSNA1 incorporation versus tubulin incorporation without SSNA1, p = 0.95; Tubulin incorporation with or without SSNA1, p = 0.004; and SSNA1 incorporation versus tubulin incorporation with SSNA1, p = 0.002 (Tukey’s multiple comparisons test). (**E)** Tubulin intensity at damage sites with or without SSNA1. p = 0.0002 (paired t-test). N = 3 independent experimental days, n = 181 damage sites for +SSNA1 condition and n = 126 damage sites for the control condition (-SSNA1). Data are binned by damage site size as indicated in legend. Data points with a border represent mean of data bin across experimental days. Lines with error bars show mean ± standard deviation.

In control conditions without SSNA1 pre-incubation, tubulin strongly localized to the sites of damage, covering 79% of the detected damage at sites smaller than 0.5 µm and more than 50% at larger sites (0.79 ± 0.02 coverage for damage site sizes < 0.5 µm, 0.59 ± 0.01 for 0.5 – 1 µm, 0.54 ± 0.15 for 1 – 3 µm), consistent with efficient tubulin-mediated lattice damage repair (Reid et al. 2017). When microtubules were preincubated with SSNA1, SSNA1 also efficiently detected and covered damage sites of all sizes (Fig. 3D, 0.75 ± 0.04 coverage for damage site sizes < 0.5 µm, 0.60 ± 0.08 for 0.5 – 1 µm, 0.64 ± 0.06 for 1 – 3 µm). However, when soluble tubulin was added after preincubation with SSNA1, tubulin localization was restricted to the edges of the damaged regions (Fig 3B). Both the degree of coverage and the intensity of the average tubulin signal were significantly lower when SSNA1 was present at damage sites than in control conditions (Fig. 3D-E, 0.64 ± 0.13 fraction of damage covered and an average of 4.0 ± 0.5 a.u. tubulin intensity in the control condition versus 0.36 ± 0.08 coverage and 2.3 ± 0.3 a.u. tubulin intensity in SSNA1 pre-incubation condition). We thus conclude that, rather than promoting tubulin-mediated lattice repair, the presence of SSNA1 limits tubulin incorporation into the sites of microtubule lattice damage.

### SSNA1 does not detect repaired microtubule damage

Our finding that the preincubation of damaged microtubules with SSNA1 restricts subsequent tubulin localization led us to question whether tubulin and SSNA1 compete for binding to microtubule damage sites. To investigate this, we modified our experimental assay to simultaneously introduce SSNA1 and tubulin to damaged microtubules (Fig. 4A). We compared tubulin localization over the same range of damage site sizes in the presence of SSNA1 to that in control conditions without the addition of SSNA1 (Fig. 4B-C). Surprisingly, we found no difference in either tubulin intensity or fraction of damage site coverage by tubulin between the control and +SSNA1 conditions (Fig. 4D-E, 0.52 ± 0.23 fraction of damage covered and an average of 6.7 ± 5.4 a.u. tubulin intensity in the control condition versus 0.50 ± 0.27 coverage and 5.4 ± 3.5 a.u. tubulin intensity when SSNA1 was added simultaneously). Thus, the presence of SSNA1 in solution does not prevent tubulin-mediated microtubule lattice repair.

**Figure 4.**
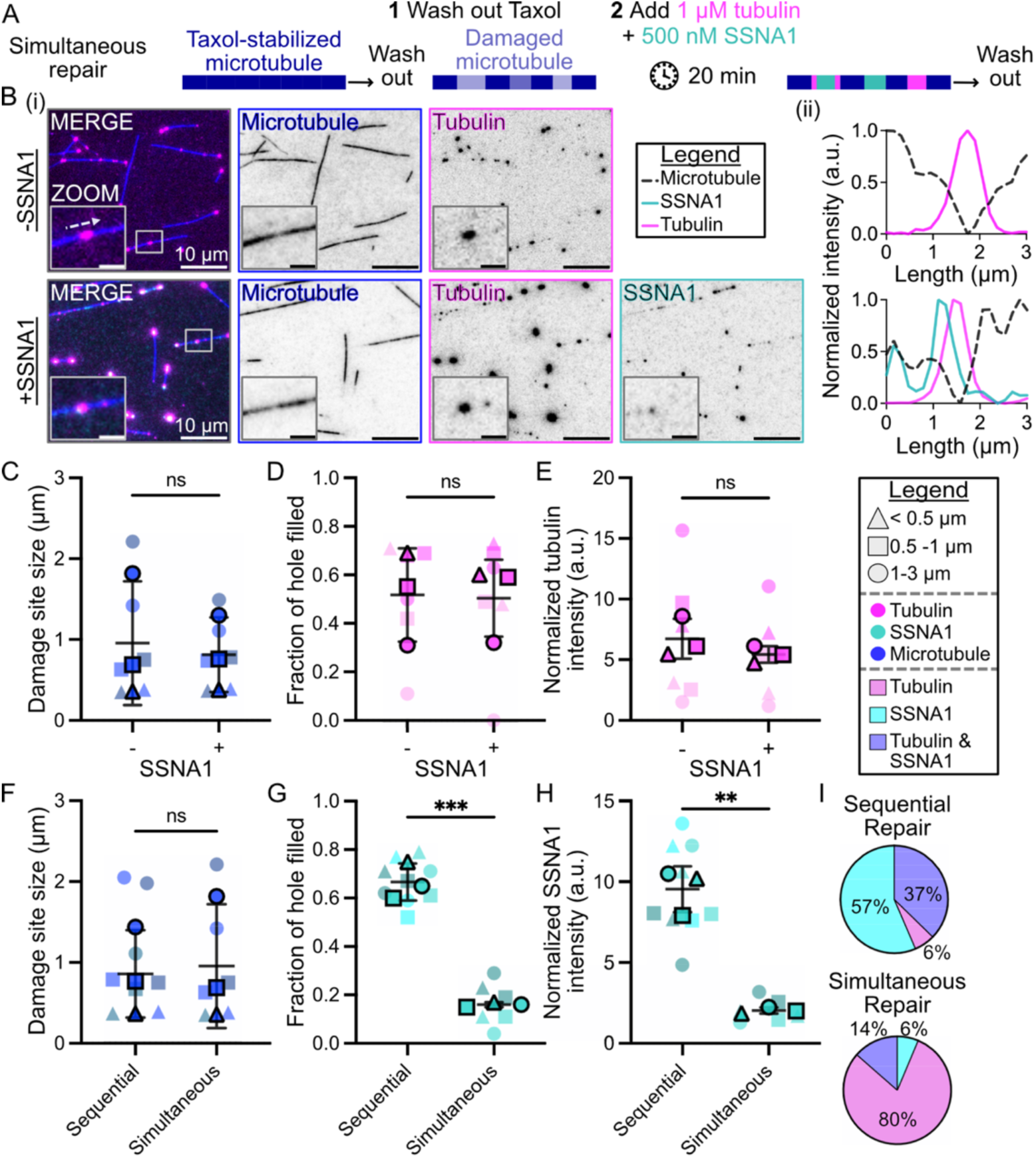
SSNA1 does not detect repaired microtubule damage. **(A)** Diagram of experimental setup. (**B)** Representative experimental images. Zoom scale bars represent 2 µm. (**C)** Distribution of damage site sizes detected during analysis in ± SSNA1 conditions. p = 0.3 (paired t-test). (**D)** Comparison of tubulin incorporation at damage sites with or without SSNA1. p = 0.8 (paired t-test). (**E)** Comparison of tubulin intensity at damage sites with or without SSNA1. p = 0.3 (paired t-test). N = 2 independent experimental days, n = 95 damage sites for +SSNA1 condition and n = 36 damage sites for control condition (-SSNA1). (**F)** Distribution of damage site sizes detected during analysis in conditions where tubulin was added either after SSNA1 (sequential) or at the same time as SSNA1 (simultaneous). p = 0.42 (paired t-test). (**G)** SSNA1 incorporation at damage sites where tubulin was added either after SSNA1 (sequential) or at the same time as SSNA1 (simultaneous). p = 0.0002 (paired t-test). (**H)** SSNA1 intensity at damage sites where tubulin was added either after SSNA1 (sequential) or at the same time as SSNA1 (simultaneous). p = 0.004 (paired t-test). n = 181 damage sites, N = 3 independent experimental days (sequential); n= 95 damage sites, N = 2 independent experimental days (simultaneous). Data are binned by damage site size as indicated in legend. Data points with a border represent mean of data bin across experimental days. Lines with error bars show mean ± standard deviation. (**I)** Comparison of protein coverage at each pixel of damage sites in sequential versus simultaneous repair conditions. n = 657 damage pixels, N = 3 independent experimental days (sequential); n = 236 damage pixels, N = 2 independent experimental days (simultaneous).

In contrast, the presence of soluble tubulin reduced SSNA1’s localization to sites of microtubule damage. Indeed, when we compared microtubule damage and repair assays where SSNA1 and tubulin were added simultaneously to those where tubulin was added following an incubation with SSNA1, we saw a significant reduction in both the degree of damage coverage by SSNA1, as well as in SSNA1’s intensity at damage sites in simultaneous addition conditions (Fig. 4F-H, 0.67 ± 0.09 fraction of damage covered and an average of 9.4 ± 2.9 a.u. SSNA1 intensity when SSNA1 was first added to damage versus 0.16 ± 0.09 coverage and 2.0 ± 0.7 a.u. SSNA1 intensity when SSNA1 and tubulin were added to damage simultaneously). Furthermore, we compared tubulin and SSNA1 co-localization at damage sites when tubulin was added sequentially after SSNA1 vs when tubulin and SSNA1 were added simultaneously to damaged microtubules. When tubulin was added sequentially after SSNA1, 94% of the pixels classified as damage were occupied by SSNA1, with 38% showing co-localization of tubulin with SSNA1 (Fig. 4I, top). In contrast, when both tubulin and SSNA1 were added simultaneously, tubulin covered 94% of damage site pixels, and the overlapped SSNA1 and tubulin colocalization was reduced to 14% (Fig. 4I, bottom). Taken together, these results suggest that SSNA1 does not detect microtubule lattice damage that has been repaired by soluble tubulin.

## Discussion

Our results demonstrate that SSNA1 functions as a direct mechanical stabilizer of the microtubule lattice by increasing bending stiffness and protecting against rupture under both motor-generated and hydrodynamic forces. SSNA1’s direct detection of microtubule lattice damage and preferential accumulation at sites of elevated mechanical strain supports a model in which SSNA1 is recruited by strain-amplified lattice defects, where it stabilizes microtubule lattice and suppresses mechanically induced failure. Previous reports found that MAP2 and tau also increase microtubule bending stiffness (Dye et al. 1993; Nishida et al. 2023). However, these MAPs bind along and reinforce undamaged microtubule lattices. Interestingly, tau binding was reported to promote microtubule lattice compaction and, unlike for SSNA1, mechanical bending of the microtubule polymer reduced tau localization (Prezel et al. 2018; Schaap et al. 2007; Siahaan et al. 2022). Microtubule acetylation, a luminal posttranslational modification of the microtubule polymer, has also been implicated in regulation of microtubule mechanics (Portran et al. 2017; Xu et al. 2017). Similar to the effects of SSNA1, acetylation protects microtubules against mechanically induced polymer rupture. However, unlike SSNA1, acetylation enhances lattice flexibility, reducing the bending stiffness of microtubules (Xu et al. 2017; Portran et al. 2017). Thus, SSNA1’s ability to specifically reinforce sites of microtubule damage and increased lattice strain provides a novel mechanism of enhancing microtubule polymer stability.

Unexpectedly, SSNA1 restricts rather than promotes tubulin incorporation at damage sites, setting it apart from known damage-recognition proteins such as CLASP, CSPP1, CLIP-170, and Abl2 (Duan et al. 2023; Van Den Berg et al. 2023; Aher et al. 2020; de Forges et al. 2016). Unlike SSNA1, all of these MAPs bind soluble tubulin and promote tubulin-mediated lattice repair (Mishima et al. 2007; Al-Bassam et al. 2010; Duan et al. 2023; Van Den Berg et al. 2023; Aher et al. 2020; de Forges et al. 2016). The observed inhibition of tubulin incorporation by SSNA1 suggests a steric mechanism, likely involving cooperative SSNA1 fibril formation across damaged lattice regions which physically occludes tubulin access. Furthermore, the inability of SSNA1 to recognize sites that have already been repaired by tubulin further indicates that SSNA1 specifically detects unique structural features of the damaged lattice. Together, these findings suggest that SSNA1 defines a previously unrecognized stabilization pathway in which microtubules are mechanically reinforced without tubulin-mediated lattice repair. Such a mechanism may be particularly important under conditions of low soluble tubulin availability or sustained mechanical load, consistent with SSNA1’s cellular localization to axonal branch points and basal bodies of cilia (Basnet et al. 2018; Lai et al. 2011; Pfannenschmid et al. 2003; Schoppmeier et al. 2005). We thus propose that preserving microtubule integrity through local mechanical lattice reinforcement provides an important complementary mode of physiologically relevant cytoskeletal stabilization.

## Acknowledgments

We are grateful for the support and resources provided by the Vanderbilt Institute of Nanoscale Science and Engineering (VINSE), where a portion of this research was conducted. We acknowledge the support from NIH R35 GM1192552 and NSF ID 2018661 grants to M. Zanic and the NSF GRFP Grant No.

1937963 to L. Richardson. The authors would also like to thank the entire Zanic lab for their assistance and support.

## Materials and Methods

### Protein expression and purification

Human SSNA1 cDNA in a bacterial expression vector with an N-terminal 6xHis tag was purchased from GeneCopoeia (Rockville, MD; product ID: Q0661). Human K560 (NM_004521.3) with a C-terminal eGFP-6His tag in a pET17b vector was purchased from Addgene (Watertown, MA; plasmid # 15219). cDNA for both constructs was transformed into Rosetta2 (DE3)pLysS cells (Novagen, Madison, WI) via heat shock. Once bacterial cultures reached an OD600 of 0.6, protein overexpression was induced with 1 mM IPTG for 18 hours at 18 C. Bacterial pellets were flash frozen in LN2. Protein expression was evaluated using SDS-PAGE.

Expression pellets were lysed for 20 minutes at 4C in 50 mM HEPES (pH 7.4), 150 mM NaCl, 1 mM TCEP, 10 mM Imidazole (pH 7.0), 0.01% (v/v) Igepal CA-630, 0.5 mg/mL chicken egg white lysozyme (Sigma-Aldrich, Saint Louis, MO), and EDTA-free protease inhibitor (Roche Diagnostics, Indianapolis, IN). Lysed cells were supplemented with 0.02% (v/v) Deoxycholate detergent (Thermo Scientific, Waltham, MA) and 0.5 M NaCl, then sonicated on ice with a Q500 Sonicator (QSonica, Newtown, CT). Crude lysate was clarified at 16,000 x g in an accuSpin Micro 17R microcentrifuge (Thermo Scientific, Waltham, MA). The supernatant was loaded onto a HisTrapFF column (Cytiva, Marlborough, MA) using an ÄKTA Pure HPLC (Cytiva, Marlborough, MA). Bound protein was washed with high salt (250 mM) for 10 column volumes and eluted with a linear gradient of 150-500 mM Imidazole solution containing 50 mM HEPES (pH 7.4), 150 mM NaCl, 1 mM TCEP, 0.01% (v/v) Igepal CA-630, 5% glycerol. Protein purity was assessed by SDS-PAGE. Amicon centrifugal spin columns (Millipore Sigma, Burlington, MA) were used to concentrate protein and buffer exchange into storage buffer with 20 mM HEPES (pH 7.4), 150 mM NaCl, 1 mM TCEP, and 10% (v/v) glycerol. Labeled SSNA1 and K560 protein fractions were further purified using size exclusion chromatography (SEC) on a Superdex 200 Increase 10/300 GL column (Cytiva, Marlborough, MA). Bovine brain tubulin was purified as previously described (Castoldi and Popov 2003).

### Protein labeling

SSNA1 protein was chemically labeled with AlexaFluor647 NHS Ester (Succinimidyl Ester) dye (Invitrogen, Carlsbad, CA; cat. A37573). Briefly, SSNA1 protein in sodium bicarbonate buffer (pH 8.25) was added to dye dissolved in dimethylsulfoxide (DMSO) at a 1:10 molar ratio (protein:dye) and rotated in a tube protected from light for 30 minutes at room temperature. Conjugated protein and excess dye were separated using a Micro Bio-Spin P6 Gel Column (BioRad, Hercules, CA; cat. 7326221). Degree of labeling was measured by nano-drop. Tubulin was labeled according to previously described methods (Hyman et al. 1991).

### TIRF microscopy imaging

All imaging was performed using a Nikon Eclipse Ti microscope with a 100×/1.49 n.a. TIRF objective (Nikon, Tokyo, Japan), Andor iXon Ultra EM-CCD (electron multiplying charge-coupled device) camera (Andor, Belfast, UK); 488, 561, and 640 nm solid-state lasers (Nikon Lu-NA); HS-625 high speed emission filter wheel (Finger Lakes Instrumentation, Lima, NY); and standard filter sets. TIRF assays were carried out as previously described (Gell et al. 2010). Briefly, chambers for imaging were created by sandwiching three thin strips of parafilm between 22 x 22 mm and 18 x 18 mm ultraclean and silanized coverslips. Microtubules were bound to the surface of the coverslip by anti-rhodamine antibody (Invitrogen, Carlsbad, CA; Cat. A-6397) and non-specific binding was blocked by incubation of pluronic F127 in the chambers. To minimize photobleaching, the Imaging Buffer consisted of 40 mM Glucose, 40 μg/ml Glucose oxidase, 16 g/ml Catalase, 0.16 mg/ml Casein, 5 mM DTT, and 50 mM KCl in 1x BRB80.

### Microtubule gliding assay

Taxol-stabilized microtubules were polymerized at 35°C from purified bovine tubulin 25% labeled with Tetramethylrhodamine (TAMRA) for 1 hour in BRB80 (80 mM PIPES, 1 mM EGTA, 1 mM MgCl2) supplemented with 1 mM of Guanosine-5’-triphosphate (GTP) and 4 mM of MgCl2. Polymerized microtubules were stabilized by dilution into a 1 mM Taxol solution (Tocris, Minneapolis, MN; Cat. 1097) and spun for 5 minutes at 30 psi in an Airfuge Air-Driven Ultracentrifuge (Beckman Coulter, Indianapolis, IN). The microtubule pellet was resuspended in 1 mM Taxol in BRB80. The supernatant was discarded and resuspended microtubules were stored away from light at room temperature and used the same day.

Coverslips were coated with anti-GFP antibody (Invitrogen, Carlsbad, CA; cat. A11122) followed by surface passivation by 1% w/v Bovine Serum Albumin (BSA; Boston BioProducts, Milford, MA; cat. P-753). 100 nM K560-eGFP was introduced to passivated coverslips for 5 minutes and then washed out with BRB80 supplemented with 1 mM Taxol (BRB80-T). Taxol-stabilized microtubules (25% labeled with TTR) were flown into the channel and incubated until the desired density was reached, as visualized by TIRF microscopy. Next, 500 nM SSNA1 protein (20% labeled with AlexaFluor647) was introduced into the channel and incubated for 20 minutes. After incubation, excess SSNA1 was washed out of solution with BRB80-T. Next, additional Taxol-stabilized microtubules were added to the channel, to serve as an internal control population not coated by SSNA1 and again washed after an appropriate density was reached. Gliding was initiated by the flow of motility buffer (50 mM Na-PIPES, 50 mM Potassium Acetate, 4 mM Magnesium Sulfate, 1 mM EGTA, 0.05p w/v BSA, pH 7 supplemented with oxygen scavenger mix (40 mM Glucose, 40 μg/ml Glucose oxidase, 16 g/ml Catalase, and 5 mM DTT), 0.16 mg/ml Casein, 50 mM KCl, 1 mM Taxol, and 1 mM Adenosine Triphosphate (ATP; ThermoScientific, Waltham, MA; cat. R0441)). Gliding microtubules and SSNA1 localization were imaged every 3 seconds for 20 minutes.

GMPCPP (Guanosine-5’-[(α,β)-methyleno]triphosphate, Jena Bioscience, Jena, Germany; cat. NU-405) stabilized microtubules were also used in gliding assays to investigate the rupture frequency of microtubule populations. These experiments were carried out in the same manner as gliding assays described above with the following exceptions. Microtubules were nucleated in the presence of GMPCPP nucleotide (25% labeled with TTR) according to standard protocols (Gell et al. 2010; Chen and Doxsey 2012). In brief, 1.5 µM tubulin was mixed with 1 mM GMPCPP and 1 mM MgCl2 in 1x BRB80 buffer overnight at 35 °C. To remove unpolymerized tubulin, microtubules were spun for 5 minutes at 30 psi in an Airfuge Air-Driven Ultracentrifuge. Microtubules were incubated with 1 µM SSNA1 (40% labeled with AlexaFluor647) for 20 minutes. Gliding microtubules were imaged every 1 second for 10 minutes.

### Microfluidic device fabrication

Photolithography and device assembly were carried out as previously described (Rogers et al. 2025). In brief, silicon wafers (University Wafer, Boston, MA) were pre-cleaned with isopropyl alcohol and exposed to oxygen plasma for five minutes using a Phantom RIE-ICP (Trion, Tempe, AZ). Cleaned wafers were baked with hexadimethylsilane (HMDS) vapors to create a hydrophobic surface for enhanced photoresist adherence. MEGAPOSIT SPR 220 7.0 positive photoresist (DuPont, Wilmington, DE) was spin-coated onto treated wafers to a thickness of 10 microns. The wafers were exposed to UV light through an in-house designed photomask. Channels designed in the photomask were developed using MICROPOSIT MF-319 (DuPont, Wilmington, DE) to remove only photoresist that had been exposed to UV light. This “master” was used to create microfluidic devices by mixing polydimethylsiloxane (PDMS) elastomer and curing agent in a 10:1 ratio and pouring the mixture over master wafers in a petri dish. After baking, cured PDMS was lifted off to expose a negative imprint of the channel design in the PDMS. Devices were then cut out from the bulk PDMS using a scalpel. Inlets and outlets for tubing were punched out using biopsy punches. Final microfluidic devices were created by bonding extruded PDMS channels to 22×22 mm glass coverslips after exposure to plasma (Harrick Plasma, Ithaca, NY).

### Microtubule bending assay

GMPCPP stabilized microtubule “seeds” were polymerized according to standard protocols (Gell et al. 2010; Chen and Doxsey 2012). In brief, 3 µM tubulin (25% TAMRA-labeled) was mixed with 1 mM GMPCPP and 1 mM MgCl2 in 1x BRB80 buffer for 1 hour at 35 °C. To remove unpolymerized tubulin, microtubules were spun for 5 minutes at 30 psi in an Airfuge Air-Driven Ultracentrifuge. Microtubule seeds were bound to the surface of a microfluidic device by anti-rhodamine antibody. Dynamic extensions were grown with 12 µM of 12% AlexaFluor 488-labeled tubulin in imaging buffer supplemented with 1 mM of GTP and allowed to polymerize for 30 minutes at 35°C. Extensions were “capped” and stabilized with 2 µM of 12% AlexaFluor 488-labeled soluble tubulin and 1 mM of GMPCPP for an additional 5 minutes. Microtubules were either coated with SSNA1 by incubating 1 µM 100% AlexaFluor 647-labeled SSNA1 for 15 minutes or immediately bent. Microtubule extensions were then bent with the imaging buffer using an OB1 flow controller system (Elveflow, Paris, France) set at 30 mbar for intervals of 10-second pulses.

### Microtubule damage assay

To induce robust and consistent microtubule lattice damage, Taxol-stabilized microtubules were added into the imaging chamber and incubated for 5 minutes to allow microtubules to bind to the surface of the coverslip via attachment to anti-rhodamine antibody and washed out with imaging buffer supplemented with 1 mM Taxol. Next, the channel was washed out with imaging buffer without Taxol. As previously reported, Taxol depletion induced damage on the microtubule lattice and gradually depolymerized coverslip-bound microtubules (Aher et al. 2020; Van Den Berg et al. 2023). These conditions were kept for approximately 5 minutes, or until visual damage appeared across the lattice. After damage was seen, 1 mM Taxol was flowed back into the channel in 1x BRB80 buffer supplemented with oxygen scavenger mix. Damaged microtubules were then incubated with varying concentrations of A647-labeled (100% labeled) human SSNA1 (500 nM) and incubated for 20 minutes while imaging SSNA1 and the microtubule wavelengths every 5 seconds. After the SSNA1 incubation, the channel was washed out with 1 mM Taxol plus oxygen scavenger mix. Next, 1 µM of A488-labeled tubulin (50% labeled) plus 1 mM GTP was flowed into the chamber and incubated for 10 minutes, imaging tubulin localization every 1 second and SSNA1 and microtubule signals every 5 intervals. After tubulin localization, the channel was washed out with imaging buffer supplemented by 1 mM Taxol and at least 3 different fields of view were imaged.

### Image Analysis

#### Microtubule segmentation

Every 8th frame (every 24 seconds of 20-minute acquisition) of each microtubule gliding time-series was used in the determination of microtubule curvature. The entire dataset was not used to limit repeating measurements of individual microtubules that had not moved out of the field of view in subsequent frames. Each of the 50 frames of gliding microtubules was input into Ilastik software (Berg et al. 2019) for segmentation using a machine learning - based approach. In brief, microtubules were individually segmented based on intensity using a user-informed machine learning module. The output of the segmentation was a binary mask of each microtubule which was then input into FIJI (ImageJ) for skeletonization through the “Skeleton” plugin. After skeletonization, the “Analyze Particles” function in FIJI (Schindelin et al. 2012) was used to create ROI’s of each segmented microtubule and the coordinates of each ROI were exported as a .txt file for further analysis.

#### Microtubule extension and curvature measurements

A custom MATLAB script was used to determine the curvature of each microtubule from its discretized coordinates. The curvature analysis was restricted to microtubule lengths between 10 and 20 microns to ensure comparison between consistent microtubule populations in each condition. First, microtubules were classified into two categories of +/- SSNA1 in the following way: Microtubule ROI’s were overlayed onto the SSNA1 channel of each image. The local background intensity near each microtubule was measured by shifting the microtubule ROI by 5 pixels in the y-direction. If the shifted ROI then overlapped another microtubule or any protein aggregation on the coverslip, the ROI was manually moved to an area devoid of other microtubules or protein aggregation while remaining close to the original microtubule ROI location. The average local background intensity for each microtubule was subtracted from the corresponding average SSNA1 intensity to circumvent any variation in signal values due to inhomogeneity of the TIRF field. Finally, microtubules were classified as +SSNA1 if their normalized SSNA1 value was over 100 arbitrary units. This microtubule classification was used to index and sort individual microtubule ROI’s and then measure curvature values for each population.

To quantify the degree of bending of each polymer, each filament was represented as an ordered set of planar coordinates along its centerline

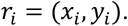

The total contour length, 𝐿_𝑐_, was computed as the sum of the Euclidean distances between consecutive points along the curve:

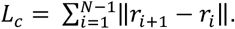

The end-to-end length, 𝐿_𝑒_, was defined as the Euclidean distance between the first and last points of the ordered filament:

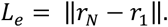

To measure the overall microtubule curvature for each population, individual microtubule’s coordinates were parameterized as arc length (s) along the polymers:

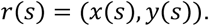

Curves were resampled at uniform arc-length spacing of 0.2 µm to obtain a discretized representation {𝑟(𝑠_𝑖_)}. To smooth any noise in data created by discretization, local tangent directions were computed using a fixed-arc baseline, Δ𝑠. At each interior position, 𝑠_𝑖_, the tangent vector was approximated by a central chord:

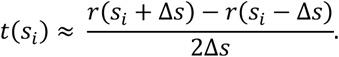

The corresponding tangent angle was defined as

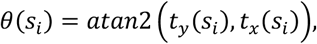

where atan2 denotes the four-quadrant inverse tangent. The sequence of tangent angles {𝜃(𝑠_𝑖_)} was unwrapped to ensure continuity along the filament.

Local bending was quantified as the absolute change in tangent angle between successive arc-length positions,

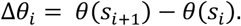

The total bending accumulated along the polymer was computed as

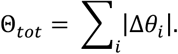

Finally, the total polymer curvature (κ) was defined by normalizing the accumulated turning by the total contour length, 𝐿_𝑐_:

### Microtubule rupture in gliding assay

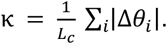

GMPCPP-stabilized microtubules were manually observed for buckling events. Each microtubule that buckled (curved) was identified and followed during gliding in the field of view. The number of buckled microtubules that rupture was compared between populations. The two populations were defined by SSNA1 intensity. SSNA1 intensity was measured for each microtubule by manually drawing ROI’s (3-pixel width lines) across each microtubule in FIJI software. These ROI’s were then overlayed onto the SSNA1 image channel to measure the average SSNA1 intensity of each ROI. Next, the ROI’s were shifted on the SSNA1 channel at least 5 pixels away from the original position to a location that didn’t overlap another microtubule. The average SSNA1 intensity along the shifted ROI was measured to give a local background measurement. Normalized SSNA1 intensities of the average intensity along the microtubule subtracted by the shifted background intensity was used to assign microtubules into populations of +/- SSNA1. Microtubules with an average normalized SSNA1 intensity over 100 a.u. were assigned to the +SSNA1 population.

### Microtubule persistence length

Microtubule persistence length was calculated using Easyworm (Lamour et al. 2014). Microtubule filaments were manually traced in the interactive MATLAB GUI. The filament centerline was represented as a parametric spline in Cartesian coordinates

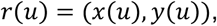

and arclength, l, was computed along the spline to obtain a contour length parameterization 𝑟(l). The total contour length of a polymer is defined as

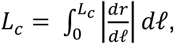

For each traced polymer, pairs of points 𝑟(𝑃_0_) and 𝑟(𝑃_0_ + 𝑙) are considered separated by a contour distance, 𝑙.

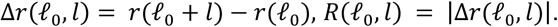

For each 𝑙, the mean-squared end-to-end distance was computed by averaging over all starting positions

𝑃_0_ along each polymer and then across filaments:

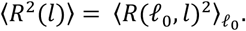

The Easyworm program then fits 〈𝑅^2^(𝑙)〉 to the worm-like chain model:

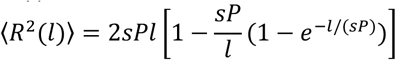

Where P, is persistence length, l, is contour length separation, and s, is a surface-equilibration parameter.

Fitting was performed by nonlinear least squares with 𝑃 as the fitted parameter.

### SSNA1 localization on bent microtubules

SSNA1 intensity was measured across different regions of bent microtubules by manually drawing ROI’s (3-pixel width lines) across each microtubule seed and extension using the tubulin fluorescent signal in FIJI software. A custom Fiji macro was used to extract intensity line profiles for all three channels from each ROI. All exported line-scan profiles were processed in MATLAB using a custom analysis script. To identify the position of the seed along each filament, the seed intensity profile was first smoothed using a moving median filter to suppress pixel-scale noise. The maximum intensity along the smoothed seed trace was identified, and the seed region was defined as the contiguous segment surrounding this maximum for which the smoothed intensity remained above a fixed fraction (50%) of the peak value. This approach robustly captured localized bright patches while excluding background or extended low-intensity regions. A minimum seed width criterion was applied to prevent single-pixel noise spikes from being classified as seeds. Next, intensity data was grouped for each microtubule into three categories of ‘seed,’ ‘next to seed,’ and ‘rest of microtubule.’ Next to seed was defined as 2 µm on either side of the seed or the total length of the extension if it is shorter than 2 µm. The rest of the microtubule was defined as any remaining microtubule lattice of the extension that was beyond 2 µm in distance. Each of these categories’ SSNA1 intensities were averaged for each microtubule and the averages for each microtubule were compared between next to seed and rest of microtubule categories using a paired t-test.

### Damage site detection

Damage sites were identified in an unbiased manner using intensity data analyzed with FIJI and a custom MATLAB script. The mean intensity of individual microtubules was measured from hand-drawn ROIs across each microtubule in FIJI and normalized by subtracting the local background intensity. Background intensity was measured by translating each microtubule ROI by approximately 5 pixels. The shifted ROI was moved further away if the 5-pixel shift resulted in the ROI overlapping another microtubule. Sites on the microtubule lattice that were 1.5 times the standard deviation below the average microtubule intensity for at least 3 contiguous pixels were considered damage sites, mathematically represented as follows:

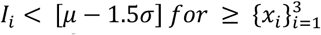

where 𝐼 is intensity, 𝜇 is the mean intensity of the microtubule, 𝑥 is distance across the damage site, and 𝜎 is the standard deviation of microtubule intensity.

### Tubulin incorporation

Tubulin and SSNA1 intensities at these damage sites were normalized by dividing raw intensity values by local background values, as determined by the same methods as above. For each pixel of detected damage sites, the pixel was considered filled by tubulin or SSNA1 if the intensity was 2 or 3 times the background intensity, respectively. This threshold was determined based on the relative labeling ratio of the proteins with the assay utilizing ∼50% labeled tubulin and 100% labeled SSNA1. Using these methods, tubulin repair efficacy was compared across experimental days binned by damage site length to prevent bias from cross-comparing different damage site sizes.

## Notes

**Competing Interest Statement:** The authors declare no conflict of interest.

### Competing Interest Statement

The authors have declared no competing interest.

